# Multiple midfrontal thetas revealed by source separation of simultaneous MEG and EEG

**DOI:** 10.1101/2020.03.11.987040

**Authors:** Marrit B. Zuure, Leighton B.N. Hinkley, Paul H.E. Tiesinga, Srikantan S. Nagarajan, Michael X Cohen

**Affiliations:** Radboud University, Donders Centre for Neuroscience; Radiology department, University of California, San Francisco; Radboud University Medical Center, Donders Centre for Medical Neuroscience

## Abstract

Theta-band (∼6 Hz) rhythmic activity within and over the medial prefrontal cortex (“midfrontal theta”) has been identified as a distinctive signature of “response conflict,” the competition between multiple actions when only one action is goal-relevant. Midfrontal theta is traditionally conceptualized and analyzed under the assumption that it is a unitary signature of conflict that can be uniquely identified at one electrode (typically FCz). Here we recorded simultaneous MEG and EEG (total of 328 sensors) in nine human subjects (7 female) and applied a feature-guided multivariate source-separation decomposition to determine whether conflict-related midfrontal theta is a unitary or multidimensional feature of the data. For each subject, a generalized eigendecomposition (GED) yielded spatial filters (components) that maximized the ratio between theta and broadband activity. Components were retained based on significance thresholding and midfrontal EEG topography. All of the subjects individually exhibited multiple (mean 5.89, SD 2.47) midfrontal components that contributed to sensor-level midfrontal theta power during the task. Component signals were temporally uncorrelated and asynchronous, suggesting that each midfrontal theta component was unique. Our findings call into question the dominant notion that midfrontal theta represents a unitary process. Instead, we suggest that midfrontal theta spans a multidimensional space, indicating multiple origins, but can manifest as a single feature at the sensor level due to signal mixing.

**Significance statement:** “Midfrontal theta” is a rhythmic electrophysiological signature of the competition between multiple response options. Midfrontal theta is traditionally considered to reflect a single process. However, this assumption could be erroneous due to “mixing” (multiple sources contributing to the activity recorded at a single electrode). We investigated the dimensionality of midfrontal theta by applying advanced multivariate analysis methods to a multimodal M/EEG dataset. We identified multiple topographically overlapping neural sources that drove response conflict-related midfrontal theta. Midfrontal theta thus reflects multiple uncorrelated signals that manifest with similar EEG scalp projections. In addition to contributing to the cognitive control literature, we demonstrate both the feasibility and the necessity of signal de-mixing to understand the neural dynamics underlying cognitive processes.

## Introduction

Oscillatory activity is driven by rhythmically active neural populations (Wang, 2010; Buzsáki and Draguhn, 2004). The temporal properties of these oscillations can inform us about the functional properties of the generating neural populations, which presumably implement the cognitive processes co-occurring with the oscillation (Womelsdorf et al., 2014). In this way, oscillatory markers of cognition can be used to understand cognition itself.

Midfrontal theta, characterized as brief theta-frequency (5-7 Hz) power increases at midfrontal EEG electrodes in humans, is well established as a marker of *response conflict*: competition between concurrently activated responses of which only one is goal-relevant (Botvinick et al., 2001). The power of the theta burst covaries with reaction time and is thought to index the level of subjective conflict (Cohen and Cavanagh, 2011). Response conflict theta is spectrally dissociable from error-related and other types of midfrontal theta (Cohen, 2014a; e.g., Kahana et al., 2001; van de Vijver et al., 2011; Cohen and van Gaal, 2014). It is non-phase-locked to stimulus or response (Cohen and Donner, 2013), suggesting that conflict enhances ongoing oscillations in the medial frontal cortex (MFC), where the signal is estimated to originate (Ridderinkhof et al., 2004). Task-relevant regions phase-lock to conflict theta (Hanslmayr et al., 2008; Cohen and Cavanagh, 2011), indicating a driving or coordinating role for conflict theta in recruiting brain networks for conflict processing. Despite extensive characterization of the signal itself, its generating neural mechanisms remain unidentified, with existing neurobiological accounts (Cohen, 2014a) being speculative at best.

Lacking mechanistic explanations, response conflict theta is assumed to be a unitary phenomenon: that is, it consists of a single signal that varies over time and correlates with the amount of conflict. Indeed, EEG and fMRI studies of response conflict theta typically capture a single source (Ridderinkhof et al., 2004), and much of the contemporary literature reflects the unitary assumption (Botvinick et al., 2001; Nigbur et al., 2011; Pastötter et al., 2013; Cavanagh and Frank, 2014; Verguts, 2017; though Töllner et al. (2017) reported two theta sources in PFC). However, this assumption may be erroneous. Because electrical signals mix linearly and instantaneously (Nunez and Srinivasan, 2006), response conflict theta may consist of multiple independent signals, but appear as a singular phenomenon at the sensor level. Similarly, the “single blob” observations typical in individual fMRI studies may hide multiple conflict-related sources, obscured by spatial normalization, smoothing, and cross-subject averaging.

In this study, we questioned the assumption that response conflict theta reflects a unidimensional neural process. We applied a feature-driven multivariate source separation method that is optimized for determining whether narrowband activity reflects the linear summation of independent sources or a single (non-linearly separable) source (de Cheveigné and Parra, 2014). This method (generalized eigendecomposition, GED) outperforms linear decomposition methods such as PCA and ICA at source separation (Nikulin et al., 2011; Cohen, 2017b; Cohen and Gulbinaite, 2017), in part because it can selectively target data features of interest.

We applied theta-targeted source separation to simultaneous MEG and EEG recorded during the “Simon task”, a task commonly used to induce response conflict (Leuthold, 2011). Our multimodal M/EEG dataset lends itself especially well to source separation, in part because of the high sensor count, and in part because MEG and EEG are sensitive to non-redundant (i.e., predominantly tangentially vs. radially oriented) neural sources. All nine subjects exhibited multiple linearly separable sources that contributed to sensor-level midfrontal theta. Inferential statistics confirmed that these sources explained significant amounts of variance in the data, and further analysis revealed that sources were temporally uncorrelated and asynchronous, implying uniqueness. Midfrontal theta thus does not appear to be a singular phenomenon, but rather a linear combination of many theta-frequency signals with overlapping topographies.

We conclude that midfrontal theta spans a multidimensional space, indicating multiple origins, and that it can manifest as a single feature at the sensor level due to signal mixing. We speculate that these dimensions reflect independent computations during conflict processing.

## Materials and Methods

### Subjects

Ten subjects (7 female) from the University of California San Francisco (UCSF) community participated in this study. The UCSF Internal Review Board (IRB) approved the study and all research was performed in accordance with UCSF IRB regulations. All subjects gave written informed consent. Inclusion criteria were the absence of psychiatric and neurological disorders, absence of substance dependence or substance abuse, and MRI safety criteria.

### Stimulus, task, and study design

Subjects performed a conflict-inducing Simon task (Figure 1A), previously described in Cohen and Ridderinkhof (2013). A colored circle was presented on the monitor, to the left or right of the fixation cross. Subjects were instructed to perform a left or right button press in response to the stimulus color while ignoring the stimulus location. Response conflict was present on trials where the target response was contralateral to the stimulus location (“incongruent trials”). Conversely, no response conflict was present on trials where the target response was ipsilateral to the stimulus location (“congruent trials”). Subjects completed 1500 trials in blocks of 60.

**Figure 1.**
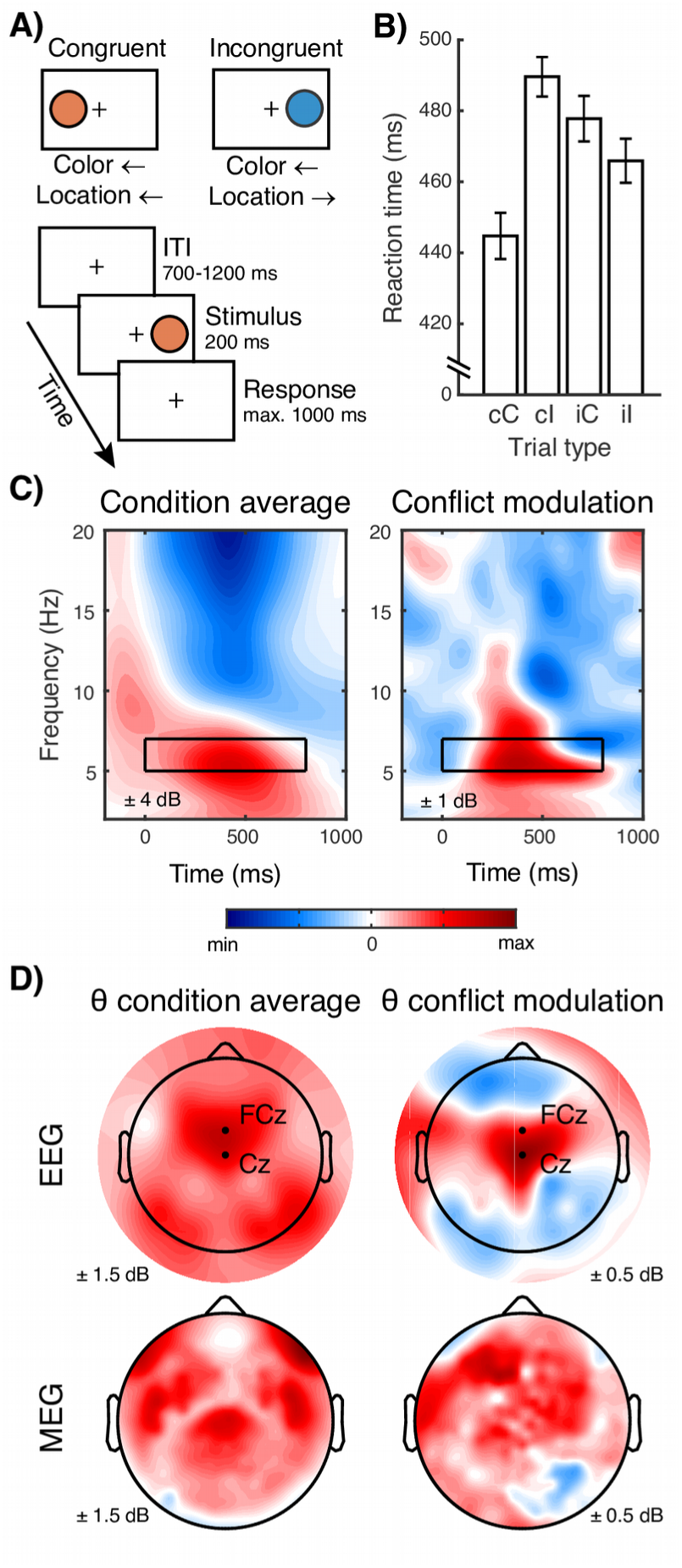
Task design, behavioral effects, and sensor-level effects. **A)** Subjects performed a Simon task with congruent (one direction cued) and incongruent, i.e., conflict-inducing (two directions cued) trials. **B)** Reaction times (± SEM) per trial type show the expected Simon effect (faster responses on congruent than incongruent trials) and expected modulation of conflict effects by previous conflict. cC = congruent following congruent, cI = incongruent following congruent, iC = congruent following incongruent, iI = incongruent following incongruent. **C)** Condition-average and conflict modulation of time-frequency power at electrode FCz. The black rectangle indicates the time-frequency window used to create the covariance matrices for the GED. **D)** Subject-averaged EEG and MEG topographies for theta (θ) power modulations.

### M/EEG recording and preprocessing

MEG data were recorded in a shielded room using a whole-head 275 axial gradiometer MEG system with third-order gradient correction (CTF MEG International Services Ltd., Coquitlam, BC, Canada) at a sampling rate of 1200 Hz. Three MEG sensors (MLF62, MLF64, MLT16) were inactive. 56 EEG electrodes were placed on the scalp according to the 10-20 system. M/EEG data were acquired under a bandpass filter of 0.001–300 Hz. Third-order gradient noise correction filters were applied to the MEG data and corrected for a direct-current offset, using CTF-provided routines.

Offline M/EEG cleaning and processing was performed with custom MATLAB scripts (MATLAB 2014a, The MathWorks, Natick, 2014) and the EEGlab toolbox (Delorme and Makeig, 2004). Data were segmented into epochs running from 1.5 s before to 2.5 s after stimulus onset. Trials were visually inspected and excessively noisy trials were rejected. Trials were classified into “congruent” and “incongruent” task conditions. As the recent history of conflict modulates reaction times (Gratton et al., 1992) and electrophysiological effects (Pastötter et al., 2013), potentially reflecting divergent cognitive processes, analyses were performed only on trials following a congruent trial (cf. Cohen and Ridderinkhof, 2013). Error trials were excluded from analysis. M/EEG data were further cleaned using independent components analysis in EEGlab. Components with clearly identifiable non-brain artifacts such as eye blinks or heartbeats were removed.

### Sensor-level time-frequency analysis

Data recorded at midfrontal EEG electrode FCz were time-frequency decomposed through trial-by-trial convolution with 40 complex Morlet wavelets. Morlet wavelets were constructed as follows (see also Cohen, 2014b):

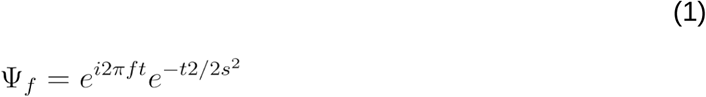

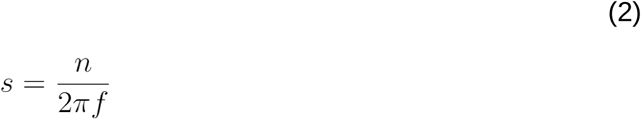

where *t* is time, *f* is frequency (logarithmically spaced from 2 to 20 Hz), *s* is the standard deviation of the Gaussian that modulates the complex sine wave, and *n* is the number of wavelet cycles (logarithmically spaced from 4 to 10). The number of cycles per wavelet governs the tradeoff between temporal and spectral precision.

Frequency-specific power was extracted as the squared amplitude of the absolute analytic signal resulting from the convolution. Power was then dB-scaled relative to a trial-averaged −500 to −100 ms baseline, 0 ms being stimulus onset. One of ten subjects exhibited uncharacteristically weak theta-frequency power at electrode FCz and was excluded from further analysis, leaving nine subjects (7 female).

### Multivariate guided source separation

The source separation method used (generalized eigendecomposition, GED; also detailed in de Cheveigné and Arzounian, 2015; Cohen, 2017c) allows for the selection of features of interest in the data. Two channel-by-channel covariance matrices **S** and **R** were constructed, based on the channels-by-time signal of interest **X**_s_ and the reference signal **X**_r_:

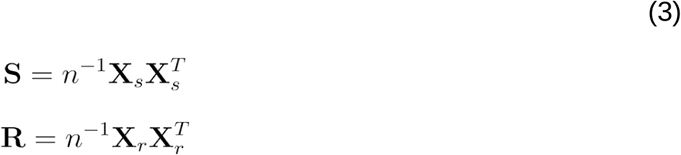

where *n* is the number of time points minus one in **X**_s_, **X**_r_. A single set of weights (in vector w) maximizing the power ratio between **S** and **R** can be found by maximizing the Rayleigh quotient (eq. 4), with scalar value λ capturing the power ratio (eq. 5).

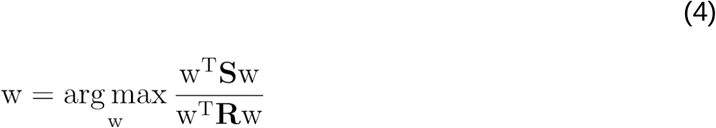

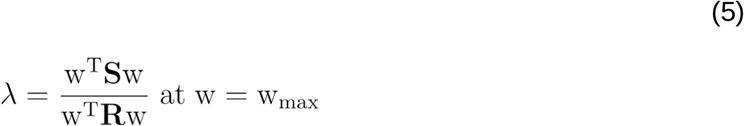

Extending the above from a single vector to a matrix computation yields the generalized eigenvalue equation in eq. 6, which, when solved, gives a matrix of weights **W** and a diagonal matrix of eigenvalues **Λ**.

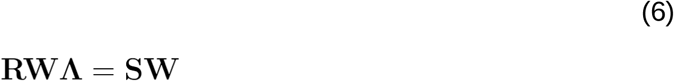

These weights (eigenvectors) capture the directions in which **S** and **R** are most separable, i.e., maximize the energy ratio between **S** and **R**. The associated eigenvalues capture the energy ratio. Notably, the eigenvectors are not constrained to be orthogonal.

We applied this eigendecomposition procedure to our data. Individual EEG trials (in μV) and MEG trials (in pT) were Z-scored to convert the data to a common scale, and combined into a single dataset per subject. Data were bandpass-filtered at theta (5–7 Hz), and signal matrix **S** was computed from a 0–800 ms time window (Figure 2). Reference matrix **R** was computed from unfiltered (broadband) data in the same time window, with 1% shrinkage regularization (eq. 7) to improve matrix separability:

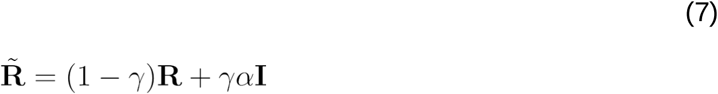

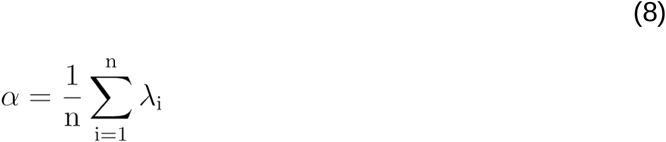

**Figure 2.**
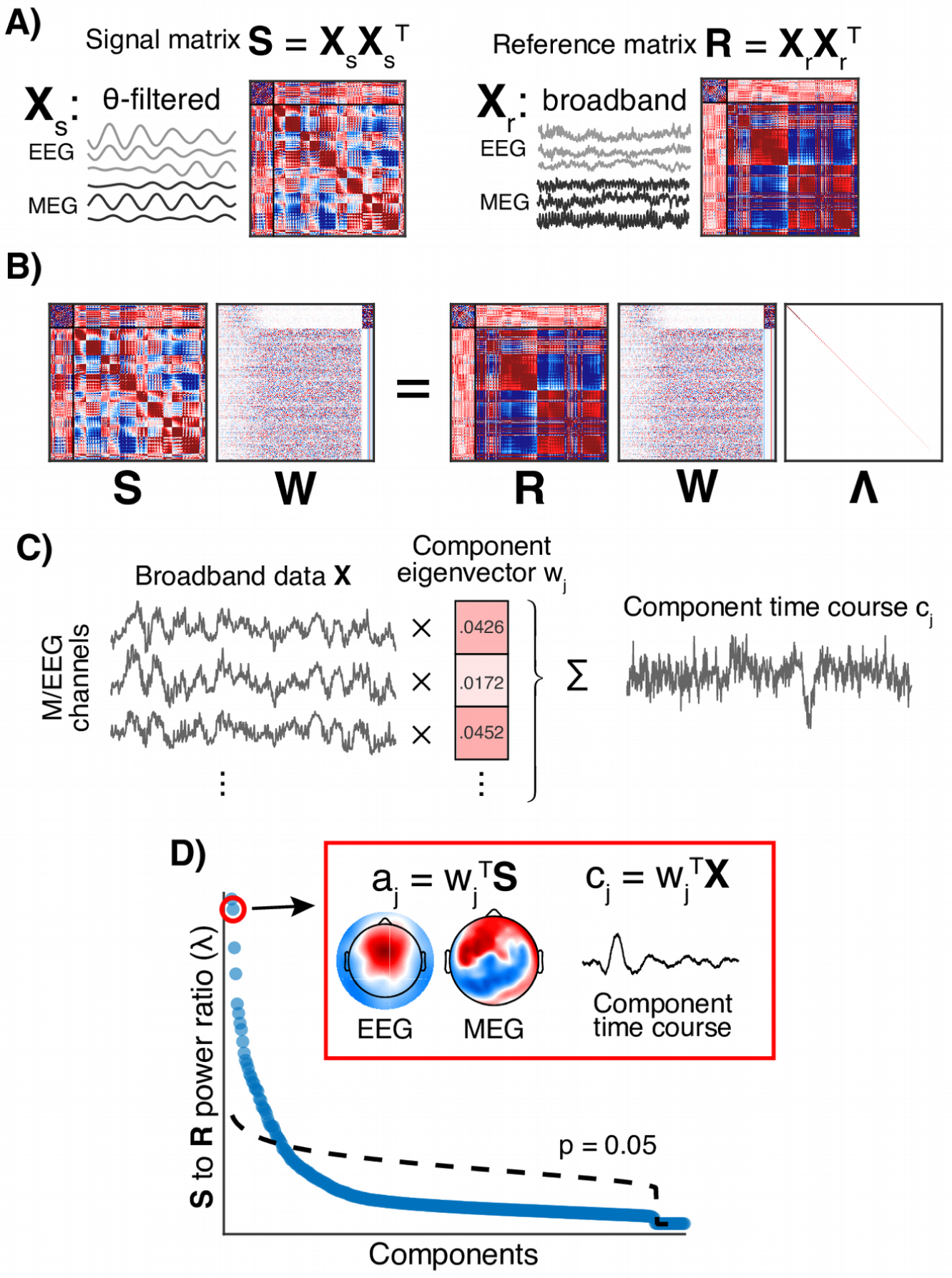
Overview of the source-separation method. A generalized eigendecomposition (GED) was performed for each subject, in order to identify statistical components contributing to theta-frequency signal. **A)** The signal matrix **S** is the covariance matrix of all M/EEG sensor signals bandpass-filtered at 5–7 Hz, from a 0–800 ms time window. The reference matrix **R** is the covariance matrix of the unfiltered sensor signals from the same time window. The first 56 rows and columns are EEG; the rest are MEG. **B)** The optimal spatial filters for theta-band activity are obtained via the GED equation. **Λ** is a diagonal matrix containing the eigenvalues, and **W** is the associated eigenvector matrix; matrices are sorted to have the largest components on the left. Note that the larger components mix EEG and MEG weightings, whereas modality-specific filters appear on the right (smaller components). **C)** The canonical time course per trial for each component is constructed as a linear combination of the time series from the 328 M/EEG sensors, using the component eigenvector as weights. **D)** Single-subject eigenspectrum, with significance cutoff (α = 0.05) determined by permutation testing. Inset shows the EEG and MEG topographies derived from the spatial filter, and the trial-averaged component time course (ERP).

where 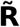 is the regularized matrix, *γ* is the degree of shrinkage (here 0.01), **I** is the identity matrix of equal size to **R**, and α is the average of the eigenvalues of **R**.

GED yielded 328 eigenvectors (“components”) for each subject, as many as there were sensors. Eigenvectors were normalized to unit length. One subject exhibited a high-energy component that was strongly driven by a single EEG electrode (F4); this electrode was removed from that subject’s dataset, and the GED was re-run. Components with repeating eigenvalues (specified as <1% difference with the previous largest eigenvalue) were excluded from analysis, as these possibly reflected inseparable components. This rejection affected only a single component with midfrontal topography. Component significance was established using permutation testing, where narrowband-filtered and broadband time series were randomly shuffled into **X**_r_ and **X**_s_ a thousand times to generate a null distribution of eigenvalues (full procedure in Hayton et al., 2004). Components explaining significant amounts of variance (α = 0.05) were retained for further analysis.

A component topography a_j_ for each given eigenvector w_j_ was computed (also shown in Figure 2D), cf. Haufe et al. (2014):

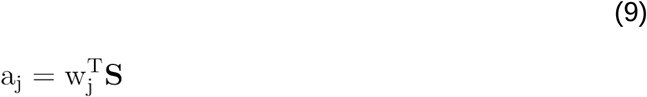

Furthermore, component time series c_j_ were constructed for each trial (also shown in Figure 2D):

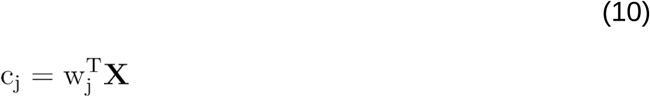

The component topography was used for topographical interpretation, and the component time series was used in all time-series analyses, described later.

### Selecting midfrontal sources

Response conflict theta manifests with stereotypical midfrontal topography on EEG recordings. To select components with midfrontal topographies, we constructed a topographical EEG template (shown in Figure 4A) that consisted of a Gaussian centered on midfrontal electrode FCz, where conflict theta power is typically maximal. Shared spatial variances (*R*^*2*^) between component EEG topographies and the midfrontal EEG template were calculated. We visually inspected the topographies and imposed a cutoff, retaining components with *R*^*2*^ > 0.5. As eigendecomposition returns eigenvectors without a canonical sign, we inverted EEG topographies so that they correlated positively with the midfrontal template, thus representing the typical conflict-related power increase over midfrontal electrodes. Furthermore, we inverted MEG topographies so that they were positive at the right lateral set of sensors. Note that sign-flipping facilitates cross-subject averaging and visual interpretation of topographies, but does not change the time-frequency response of the component time series.

**Figure 3.**
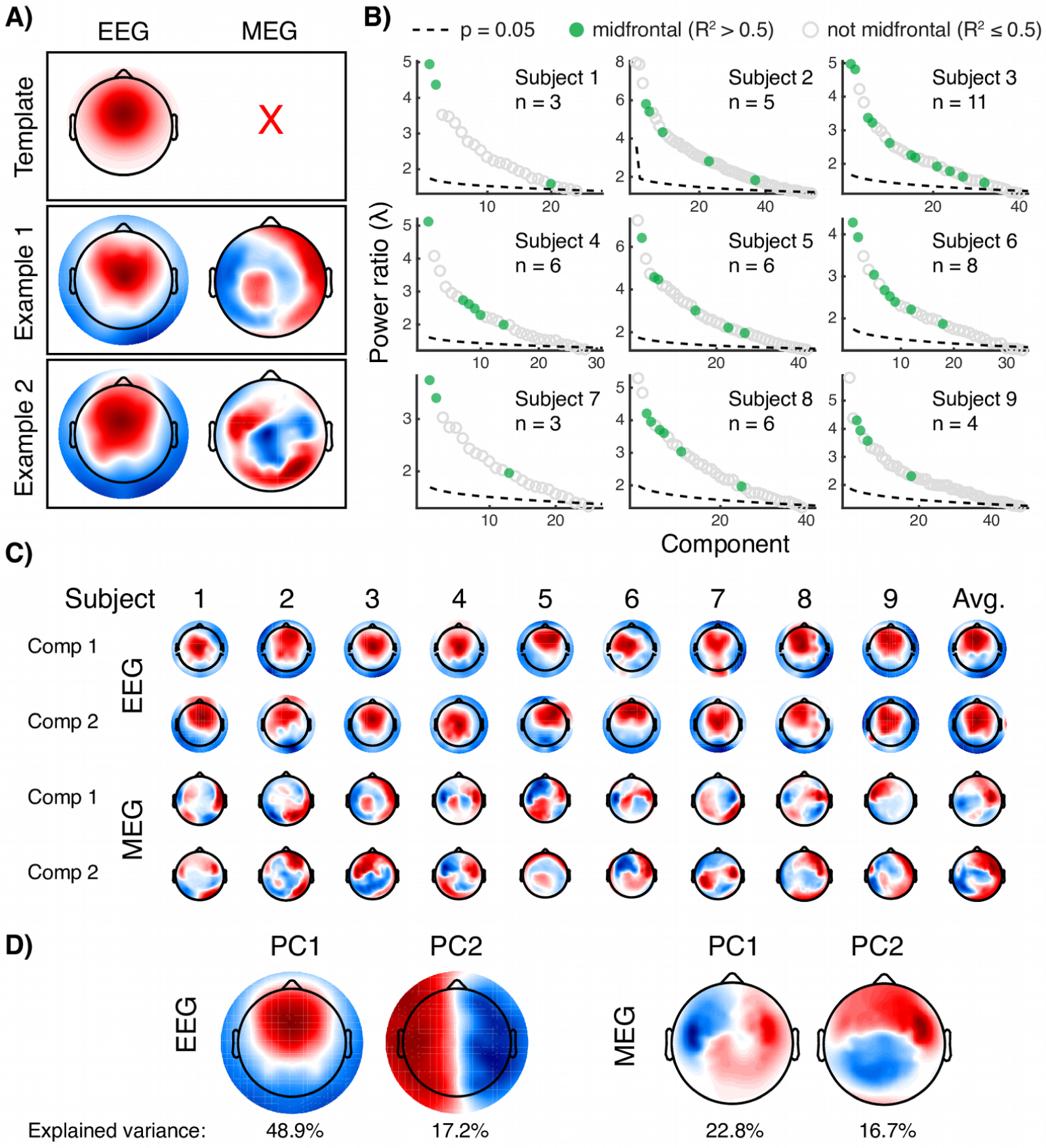
Procedure for selecting midfrontal theta components based on eigenvalue statistical significance and EEG topography template matching. **A)** Top: the EEG template topography used to select midfrontal components. Middle and bottom: components selected using this template can have diverse MEG topographies. **B)** Components sufficiently (R^2^ > 0.5) correlated with the template were retained (green). “n” indicates the number of components retained per subject. Most subjects had at least two midfrontal components in their top 5 theta components, indicating that our conclusion that midfrontal theta is driven by more than one component is not dependent on the particulars of significance thresholding. Note that components were selected based on theta energy (reflected in the power ratio) and midfrontal EEG topography; possible modulation by experiment condition was ignored during selection. **C)** The two top midfrontal components (explaining most variance) per subject exhibited homogeneous EEG topographies and heterogeneous MEG topographies. **D)** Principal component analyses individually applied to the EEG and MEG topographies captured salient spatial features across all components.

**Figure 4.**
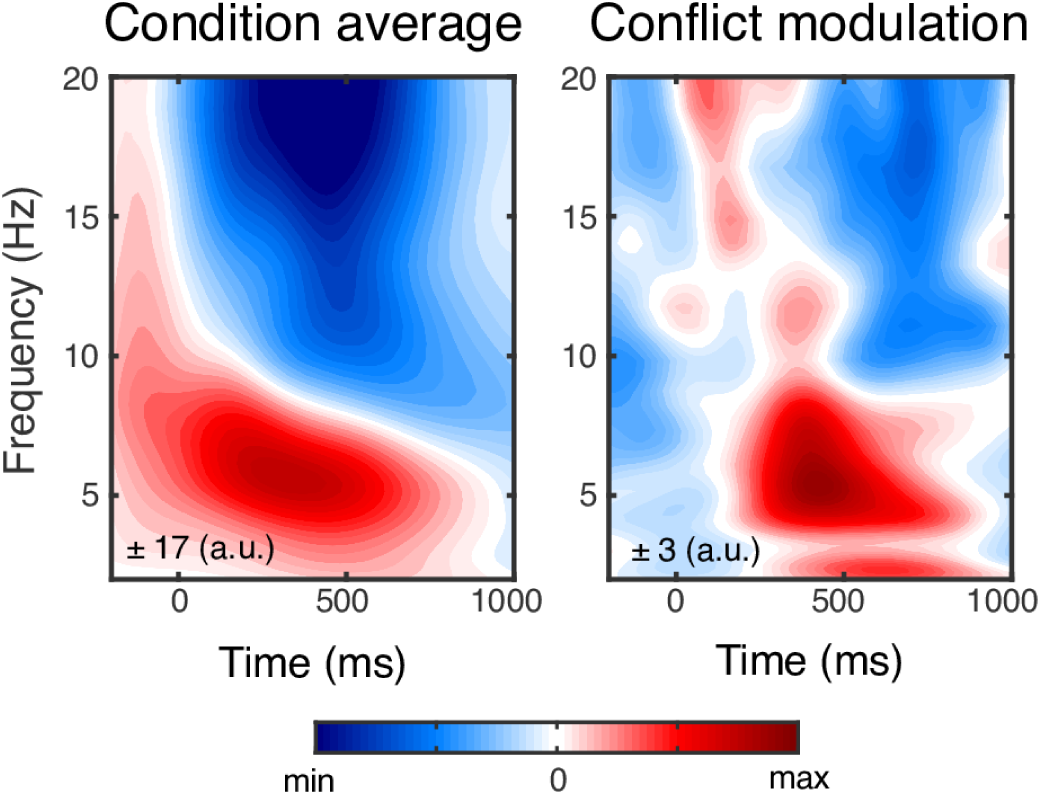
Average midfrontal component time-frequency decompositions appear qualitatively similar to those from electrode FCz (in Figure 1C). Components were weighted by their relative eigenvalue (relative to all within-subject midfrontal components), as a measure of their contribution to the overall signal.

### Evaluating source similarity

To address the possibility that GED artificially split a single source into multiple components, we evaluated components on three measures of signal similarity. In preparation, theta power (relative to a −500 to −100 ms baseline) and phase were extracted from the component time series, through convolution with a 6 Hz complex Morlet wavelet. For the first measure, pairwise component synchrony was computed as the mean length of the phase angle difference vector at each time point, and mean synchrony over −500 to 900 ms was extracted. This measure captured the component phase similarity. Synchrony between adjacent EEG electrodes Pz and POz (where synchronization was expected to be inflated by volume conduction) was computed for comparison. Second, the mean pairwise correlation between component theta power time courses was computed, capturing the similarity of within-trial power fluctuations. Third, the pairwise correlation between mean component theta power over 0–800 ms was computed, capturing the similarity of power fluctuations across trials.

Another possibility is that a single midfrontal theta source was split into two components because of noise. To investigate this possibility, we determined how much noise needed to be added to two copies of a single spatial filter in order to generate synchronization of a similar magnitude as two empirical spatial filters. This was a three-step process. First, we selected a spatial filter (the eigenvector w from eq. 4), made a copy of this filter, and then added unique white noise to each copy (noise standard deviation, SD_noise_, ranging from 0 to 10 times the eigenvector SD, SD_eig_, in steps of 0.1; 50 copies per noise level). Second, theta power time courses were derived from these noisy spatial filters. Third, the expected distribution of pairwise correlations between time courses was computed for each of the noise levels (a total of ½ × 50 × 50 - 50 = 1200 unique pairwise “noisy self-correlations” per noise level). One-tailed T-tests (α = 0.05, no multiple comparisons correction) between the correlation distributions determined the noise level for which the generated self-correlations were no longer statistically distinguishable (for at least one participant) from the empirical cross-component correlations.

### Component-level time-frequency analysis

The retained component time series were time-frequency decomposed through trial-by-trial convolution with 40 complex Morlet wavelets, as previously described for data from sensor FCz. The average time-frequency spectrum across components was computed. To accurately represent each component’s contribution to the overall energy, each component was weighted by the variance it explained among the retained components per subject.

### Computing Granger causality

To attempt to identify a hierarchy in how components interacted over time, we computed windowed Granger causality (GC) on component time series (Figure 7A) using the MATLAB MVGC toolbox (Barnett and Seth, 2014). The ERP over trials was subtracted from each trial to limit the influence of nonstationarities. Data were downsampled from 1200 to 300 Hz. Model order was estimated per subject, on all trials, over 0–800 ms, using Bayes’ information criterion. We selected the maximum order across subjects (6 time steps, equaling 19.2 ms) for all further GC calculations. Next, trial data were segmented into 200 ms windows with 50 ms step size. Conditional GC between each pair of components was computed over each time window:

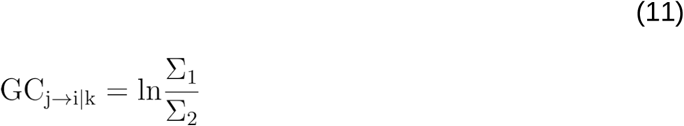

where *j* is the driving component, *i* is the receiving component, and *k* is the group of all other components, on which *j*→*i* is conditional. Σ_1_ is the error term (variance of the residual) of the autoregressive model fit to the recent history (constrained by the order parameter) of *i*, with linear terms for the influence of the components in *k*. Σ_2_ is the error term for the same model with the addition of a linear term for the influence of *j*. Thus, if the prediction for *i* improves with incorporating the past of *j*, the GC value exceeds 0. Averaging over “outgoing” and “incoming” GC values yielded a “driving mass” and “receiving mass” value for each component within each time window:

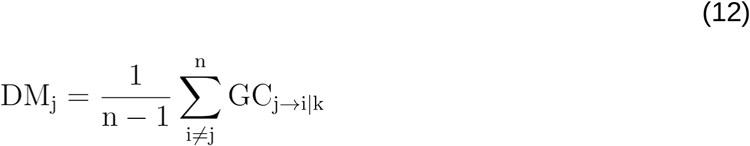

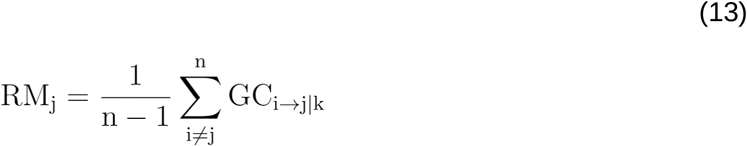

where *n* is the number of components for that subject. Driving and receiving mass were computed separately for incongruent trials and congruent trials, with condition-average GC being the average of these two.

**Figure 5.**
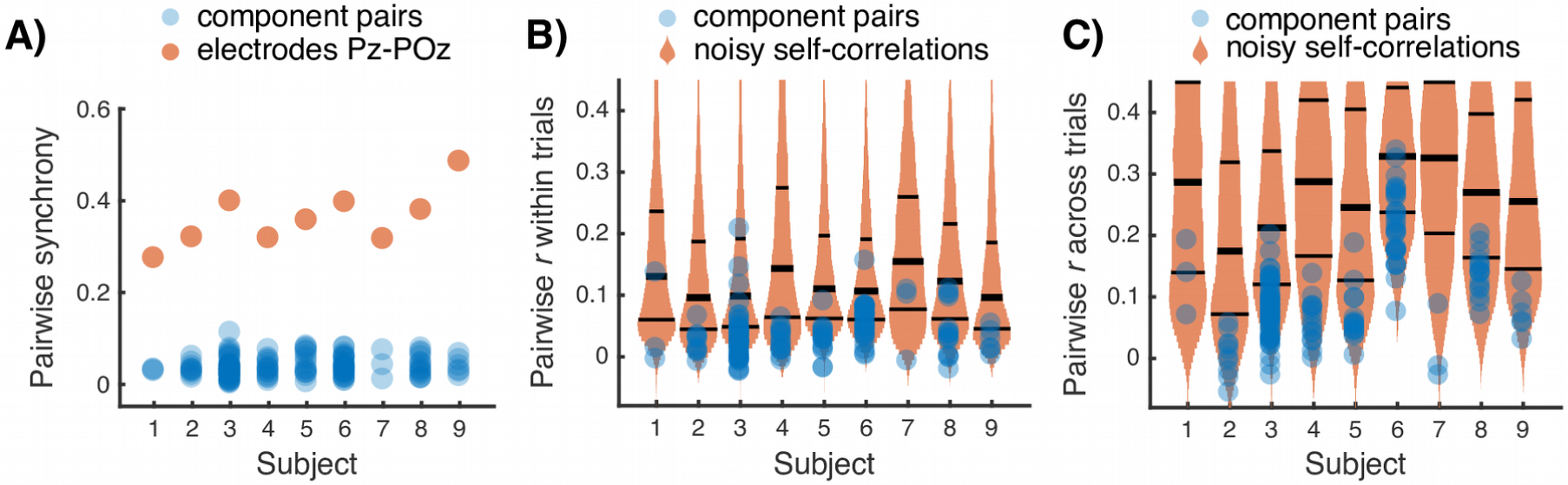
Several control analyses provided evidence that the components reflected unique data features, as opposed to single theta sources artificially split into multiple components. **A)** Pairwise synchrony across components (calculated as the mean length across trials of the phase difference vector) was generally weak. For comparison, orange dots show synchronization between Pz and POz, where the inter-electrode synchronization can be expected to be inflated by volume conduction. **B)** Trial-by-trial correlations between theta power time series for different components were generally low (blue dots). The orange violin plots indicate distributions of correlations between copies of the same component but with white noise added. At SD_noise_ = 1.9 × SD_eig_, generated correlations first became statistically indistinguishable (p > 0.05 for at least one subject) from the empirical pairwise component correlations. This suggests that the GED-identified components were unlikely to originate from shared theta sources in the presence of only modest levels of noise. Horizontal lines in the violin plots indicate mean ± SD. **C)** Similar to B, but for mean theta power per trial, and with the orange violin plots indicating the distribution of correlations for SD_noise_ = 1.5 × SD_eig_ (at which generated and empirical correlations first became indistinguishable).

**Figure 6.**
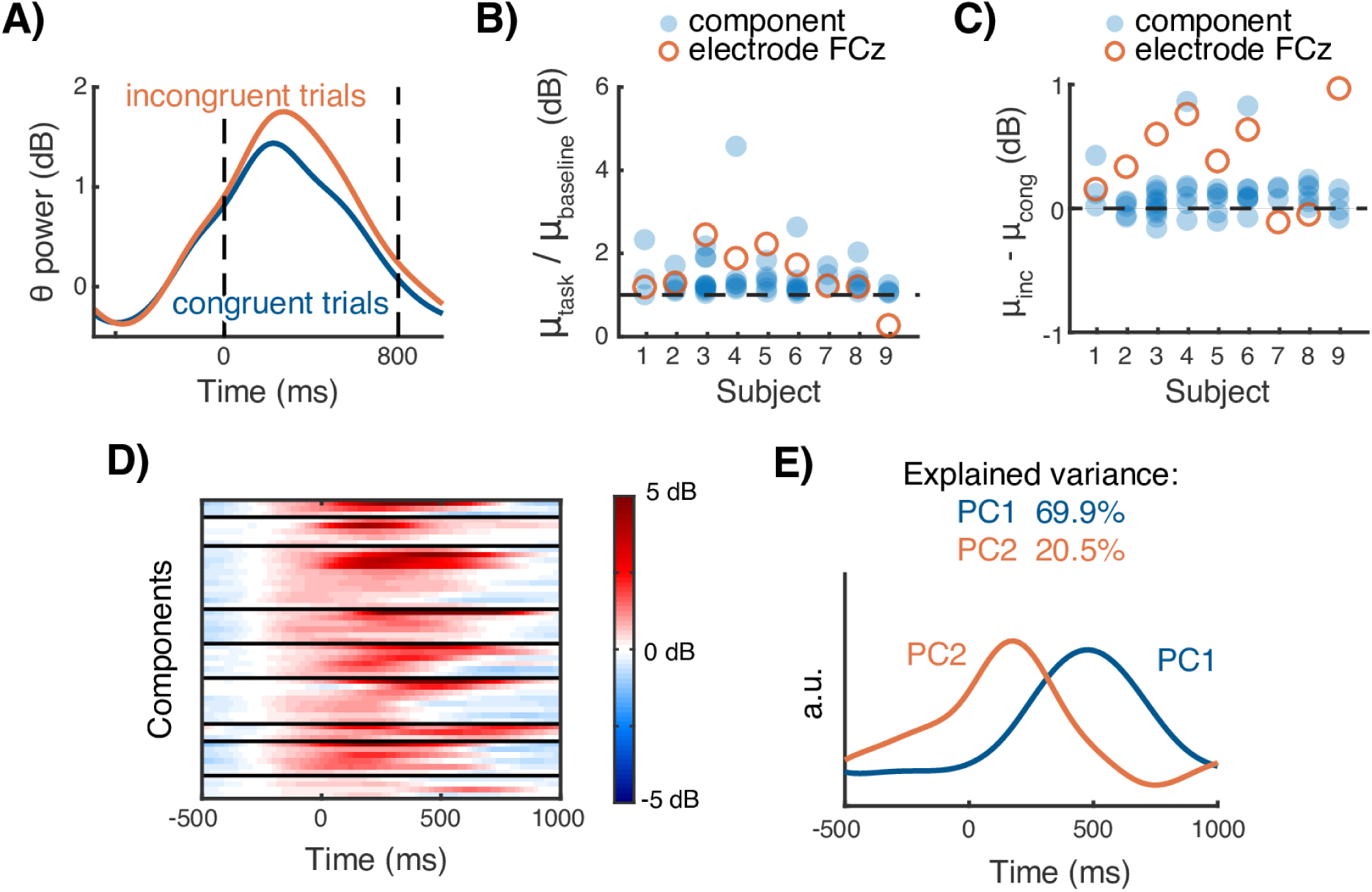
Characterization of component theta (θ) power dynamics over time and conflict conditions. **A)** Component-averaged theta power per condition. **B)** Mean theta power modulation during the task (0–800 ms) relative to baseline (−500 to −100 ms) for each component (blue dots) and electrode FCz (orange circles) for each subject. **C)** Similar to B, but showing theta power modulation during incongruent compared to congruent trials. **D)** Components exhibited variability in theta power over time. Black lines separate components originating from different subjects. **E)** PCA revealed salient temporal features of the theta power time series in panel D. The top PC reflects a later theta power increase, explaining most of the variance, and the second PC reflects an earlier theta power increase.

**Figure 7.**
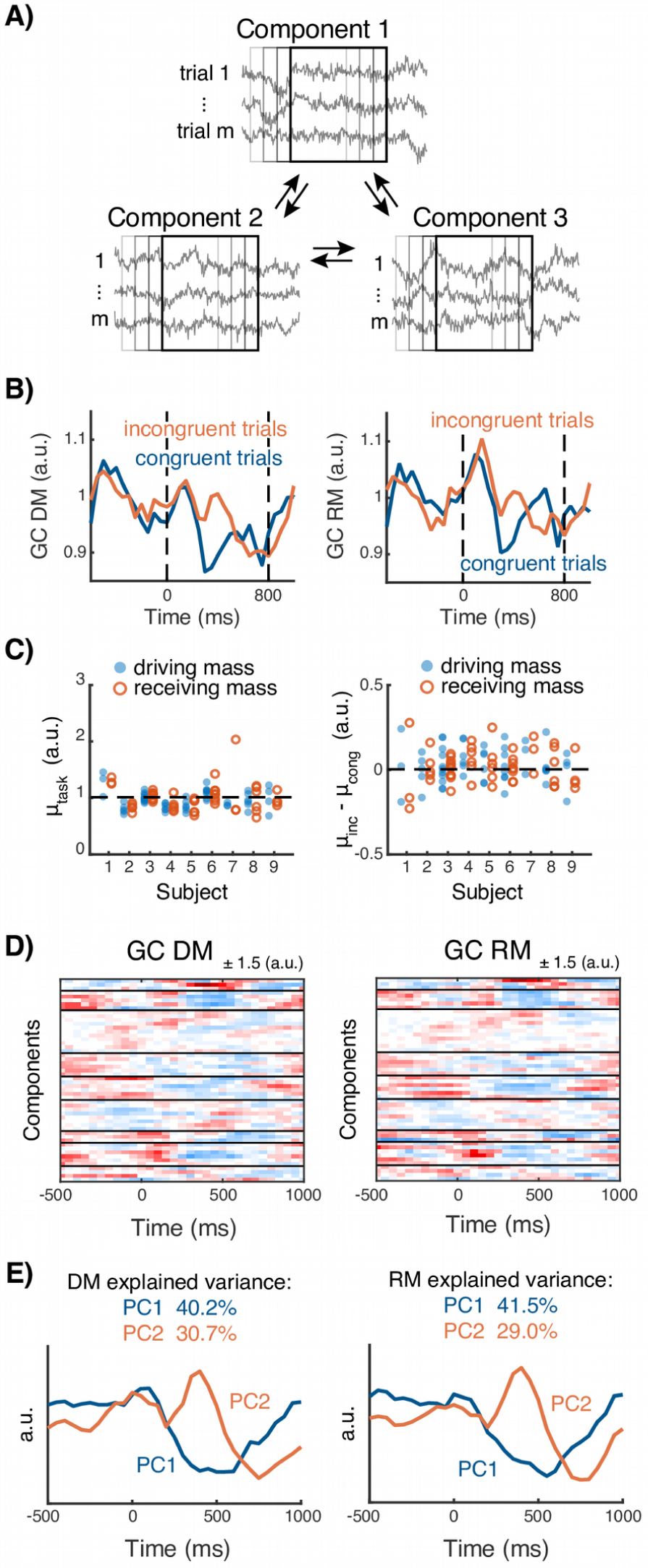
Granger causality (GC) analyses show no evidence for one component driving others, suggesting the absence of a “CEO-like” hierarchy. **A)** Conditional GC among all components per subject (varying from 3 to 11 per subject) was computed across 200 ms windows in 50 ms steps. Procedure is illustrated for three components. **B)** GC time courses, normalized to a −500 to −100 ms baseline. **C)** Average GC values per component, for baseline-normalized condition average (μ_task_) and conflict modulation (μ_inc_-μ_cong_). **D)** Time courses for DM and RM for all components (black lines demarcate subjects). **E)** PCAs on the GC time series revealed a slower decrease and an earlier and more phasic increase.

### Deriving task and conflict modulation

For each component, theta power modulation during the task was computed as mean theta power on all trials (normalized relative to a −500 to −100 ms baseline) over 0–800 ms. This yielded a ratio, i.e., a modulation value of 1 indicated no change in theta power compared to baseline. Conflict-related modulation was computed as mean theta power modulation on incongruent trials minus modulation on congruent trials. These values were also computed at midfrontal electrode FCz, for comparison purposes. One component was rejected for being strongly negatively task-modulated, leaving 52 components. Component-specific Granger causality modulation was computed in the same way as theta power modulation: once across all task conditions and once per condition, though computed over driving and receiving mass instead of theta power.

### Extracting temporal and spatial component features

We applied principal component analyses (PCAs) to identify temporal and spatial features of the component time courses and topographies that were consistent across subjects. Separate PCAs were applied to EEG topographies, MEG topographies, theta power time courses, and GC time courses. The number of PCs for each data feature was determined by visually identifying the “elbow” in the plot of the eigenvalues (i.e., the point where additional explained variance per PC sharply decreases) and retaining the PCs before that point. This approach suggested two PCs for each data feature. As PCA returns PCs without a canonical sign, we inverted the time course PCs and topography PCs so that they were positively correlated with the majority of original time courses and topographies.

### Statistical analysis

Statistical analyses were performed using MATLAB (MATLAB 2014a, The MathWorks, Natick, 2014). The significance level was set at α = 0.05. Two-sample differences were compared using one-way or two-way T-tests (described in results), with Benjamini-Hochberg (BH) step-up multiple comparisons correction (Benjamini and Hochberg, 1995) applied when testing related hypotheses. Behavioral results were compared using a two-way ANOVA to identify interactions, with BH multiple comparisons correction applied to pre-chosen contrasts. For GC marginals, tested at each time window and for each pairwise combination of components, we applied false discovery rate (FDR) multiple comparisons correction as implemented in the MVGC toolbox (Barnett and Seth, 2014). No multiple comparisons corrections were applied when determining at which noise level the randomized and empirical component correlation coefficients became indistinguishable (Figure 5). This lack of MC correction is intentional so that the identified noise level represents a lower bound. Component significance was determined by creating a null distribution using permutation testing, with a cutoff for retention set at *p* > 0.05. Correlation coefficients were computed as Pearson’s *r*. Descriptive statistics are always given as mean ± SD.

### Code and data accessibility

The custom-written MATLAB analysis code will be made available upon publication on GitHub at http://www.github.com/marrit-git/MEEG-multiple-theta-sources. MEEG data will be made available at https://data.donders.ru.nl.

## Results

### Behavioral results

We replicated the typical congruency sequence effect (a.k.a. the “Gratton effect”) of previous trial congruence affecting performance on the next trial (Gratton et al., 1992). Following congruent trials, accuracy was 94.7% ± 3.1% on congruent and 86.5% ± 7.1% on incongruent trials, whereas following incongruent trials, accuracy was 91.2% ± 3.7% on congruent and 92.0% ± 6.0% on incongruent trials (interaction between previous and current congruency *p* = 0.015, *F*(1,32) = 6.64, ANOVA). This effect was also observed in reaction times (Figure 1B): subjects responded faster on incongruent trials following incongruent (465.9 ± 22.5 ms) than congruent (489.6 ± 26.5 ms) trials, and responded slower on congruent trials following incongruent (477.8 ± 31.6 ms) than congruent (444.7 ± 35.8 ms) trials (interaction *p* = 0.007, *F*(1,32) = 8.30, ANOVA). As mentioned in the Methods section, we focused the MEEG analyses on trials following congruent trials, in order to maximize the conflict effect.

### Sensor-level analyses

Data recorded at midfrontal EEG electrode FCz were time-frequency (TF) decomposed through complex Morlet wavelet convolution. This decomposition served two purposes. The first was to confirm that task and conflict effects were present in the sensor-level data, rather than being artificially induced by the generalized eigenvalue decomposition (GED). The second was to guide the feature selection for the GED.

The average FCz time-frequency spectra (Figure 1C) exhibited a task-related theta power increase, as well as conflict modulation in the theta band. Power increased predominantly within the 4–7 Hz frequency range. This observation validated our choice of bandpass frequency filter (5–7 Hz, chosen for a priori expectations of theta manifesting around 6 Hz) for the GED signal matrix. The condition-average theta power increase spanned 0–800 ms relative to stimulus onset, guiding our selection of this time window to construct the GED signal and reference matrices.

Condition-averaged theta power (at 0–800 ms relative to a −500 to −100 ms baseline) was observed at midfrontal and lateral occipital locations in EEG, and at midfrontal and frontolateral locations in MEG (Figure 1D). Conflict-modulated theta power was maximal at midfrontal locations in EEG and was broadly midfrontally distributed in MEG.

Theta power at FCz was significantly greater during the task (1.38 ± 0.68 dB) than during baseline (*p* = 0.0001, one-sided *t*(8) = 6.08). However, FCz theta power was not significantly greater on incongruent (1.46 ± 0.78 dB) than congruent trials in the window of 0-800 ms (1.28 ± 0.65 dB; *p* = 0.14, one-sided *t*(8) = 1.15; all Benjamini-Hochberg corrected).

### Identification of midfrontal theta components

Custom spatial filters were created using generalized eigendecomposition (GED), to identify components in the data that maximally separated theta-band activity from broadband data (procedure illustrated in Figure 2). A permutation testing-determined significance threshold (α = 0.05) was applied to the resulting 328 components per participant (Figure 2D). As shown in Figure 3, each subject exhibited between 21 and 46 significant theta components (average 33.89 ± 9.47). A subset of these (5.89 ± 2.47 per subject; minimum 3, maximum 11) was midfrontal, defined as shared spatial variance *R*^2^ > 0.5 between the component’s EEG topography and the midfrontal template shown in Figure 3A. By virtue of the template-based selection method, component EEG topographies were homogeneous. However, the associated MEG topographies were highly variable (Figure 3C). This variability is also reflected in the salient topographical features identified by applying PCAs to the EEG and MEG topographies (Figure 3D): a single PC explained 48.9% of the variance in the EEG topographies, whereas the top two MEG PCs together captured 39.5% of the variance.

Averaged time-frequency decompositions of the retained components (Figure 4) were similar to those at electrode FCz (Figure 1C). This similarity suggests that this subset of the data retained the salient data features, and that applying GED had no distorting effects.

### Possible alternative accounts for multiple theta components

To rule out the possibility that the GED artificially split out a single signal source into multiple components, we computed three measures of component similarity, reasoning that components representing the same source should have highly similar time courses.

The first measure, pairwise synchrony at 6 Hz, was generally low (Figure 5A; 0.044 ± 0.005) in comparison to synchronization between EEG electrodes Pz and POz, which will be artificially inflated due to volume conduction contamination. These electrodes were significantly more synchronous than any pair of components (0.360 ± 0.062, *p* < 0.001, two-sided *t*(16) = 15.2). As such, components did not exhibit high similarity in terms of phase.

The second measure, trial-averaged pairwise correlations between theta power time courses, was also generally low (Figure 5B; Pearson’s *r* = 0.04 ± 0.02), indicating low similarity of within-trial power fluctuations. The third measure, pairwise correlations between mean theta power on each trial, was likewise low (Figure 5C; Pearson’s *r* = 0.11 ± 0.08).

Another potential explanation for observing multiple theta components is that noisy data caused a single real source to be split into multiple apparent sources. To determine the feasibility of this explanation, we created two noisy copies of each identified spatial filter, computed their component time series, and correlated these with each other. We then increased the amount of noise until the noise-driven time series correlations were no longer statistically distinguishable from the empirical time series correlations. We found that a noise factor of SD_noise_ = 1.9 × SD_eig_ for within-trial correlations, and SD_noise_ = 1.5 × SD_eig_ for cross-trial correlations, was required to reach this point (lower bound on noise; reported SD_noise_ measured at the first occurrence across subjects of *p* > 0.05 between generated and empirical correlation distributions; one-tailed *t* < 1.55 within-trial, one-tailed *t* < 1.52 cross-trial, varying df; no multiple comparisons correction). In other words, if components comprised noisy representations of the same source, the noise would have to exceed the signal by around two standard deviations to generate comparably low pairwise correlations.

Finally, we tested whether multiple statistical sources may have arisen from a single real source that changed over time (e.g., with head movement or changes in signal quality). We reasoned that this account would predict negatively correlated cross-trial power fluctuations. Pairwise correlations across trials were not significantly negative (*p* > 0.99, one-sided *t*(8) = 3.93 against μ = 0), suggesting that components were stable over time.

### Task and conflict modulation of theta power

To characterize the nature of these components, we extracted the theta power time courses from the component time series. We then quantified how theta power changed during the task (relative to baseline) and how it changed on incongruent vs. congruent trials. Results are shown in Figure 6. Component theta power was significantly greater during the task (1.04 ± 1.13 dB) than during baseline (*p* < 0.0001, one-sided *t*(51) = 6.67). Component theta power was significantly greater on incongruent trials (1.10 ± 1.19 dB) than on congruent trials (0.97 ± 1.09 dB, *p* = 0.0012, one-sided *t*(51) = 3.20; all Benjamini-Hochberg corrected).

Because the GED identified linearly separable components, simple averaging may fail to identify meaningful cross-component functional variability. We therefore applied a PCA to the component’s theta power time courses (Figure 6D), which identified a later peak (Figure 6E, PC1; peaking around 480 ms poststimulus) and an earlier peak (PC2; peaking around 180 ms poststimulus) in theta power. Pre-stimulus effects are attributable to temporal smoothing from the wavelet convolution.

### Granger causality

Response conflict-related midfrontal theta is classically considered to be a phenomenon with a singular origin. While our previous results suggest that this is not the case, we considered the related possibility that activity from one of the identified components dominated the other components. To address this, we computed Granger causality (GC) over time between components, quantifying the extent to which component time series predicted one another. All-to-one connectivity (“receiving mass”, RM) and one-to-all connectivity (“driving mass”, DM) were determined for each component. DM and RM were significant (*p* < 0.05 for all components and at all time points, chi-squared test, FDR multiple comparisons corrected), suggesting that components exchanged information. Baseline-normalized time courses are shown in Figure 7B.

We first investigated whether this information exchange was modulated by task conditions. Average component DM did not significantly differ between the task (2.43 ± 1.12, all GC values in this section × 10^−3^) and baseline (2.59 ± 1.24; *p* = 0.014, Benjamini-Hochberg-corrected α = 0.0125, two-sided *t*(51) = 2.53). The same was true for RM (task 2.39 ± 1.11, baseline 2.55 ± 1.30, *p* = 0.08, two-sided *t*(51) = 1.79). Similarly, component DM did not significantly differ between incongruent (2.46 ± 1.24) and congruent trials (2.39 ± 1.16; *p* = 0.11, two-sided *t*(51) = 1.60), and the same was true for RM (incongruent 2.42 ± 1.18, congruent 2.37 ± 1.05, *p* = 0.16, two-sided *t*(51) = 1.42; all Benjamini-Hochberg corrected). Information transfer thus remained stable on average, regardless of task features.

Next, we determined how information transfer was distributed across individual components. If a single component per subject drove the other components, we would expect to observe (1) that one component especially strongly predicts information in the other component time series, leading to increased DM for that component, and (2) that one component is especially weakly predicted by the other components, leading to decreased RM for that component. The actual per-component DM and RM, shown in Figure 7C, do not match either of these expectations. Additionally, DM and RM time series per component (Figure 7D) were highly correlated (*r* = 0.79, *p* < 0.001), suggesting strong symmetry in components’ driving and receiving tendencies.

PCAs applied to the DM and RM time courses (shown in Figure 7D) revealed salient temporal features (Figures 7E). The first PC for both DM and RM captured a slow decrease in GC. The second PC captured a sharper increase, followed by a decrease, of GC.

## Discussion

In this study, we tested the assumption that midfrontal conflict theta is a unidimensional phenomenon (Cavanagh et al., 2012; Cavanagh and Frank, 2014; Cohen, 2014a). We used feature-guided source separation to attempt to identify multivariate components contributing to midfrontal theta. All subjects exhibited multiple (5.89 ± 2.47) midfrontal components that contributed to increases in theta power during a response conflict task. The identified midfrontal components explained significant amounts of variance in the data, reflected unique data features, and remained stable over time. Thus, conflict-related midfrontal theta exists in a high-dimensional signal space, whose basis vectors are sufficiently similar that they appear as a single dimension without careful inspection and multivariate analysis methods (Figure 8). This multidimensionality suggests that the midfrontal conflict theta signal consists of the aggregate activity of many, diverse, theta generators.

**Figure 8.**
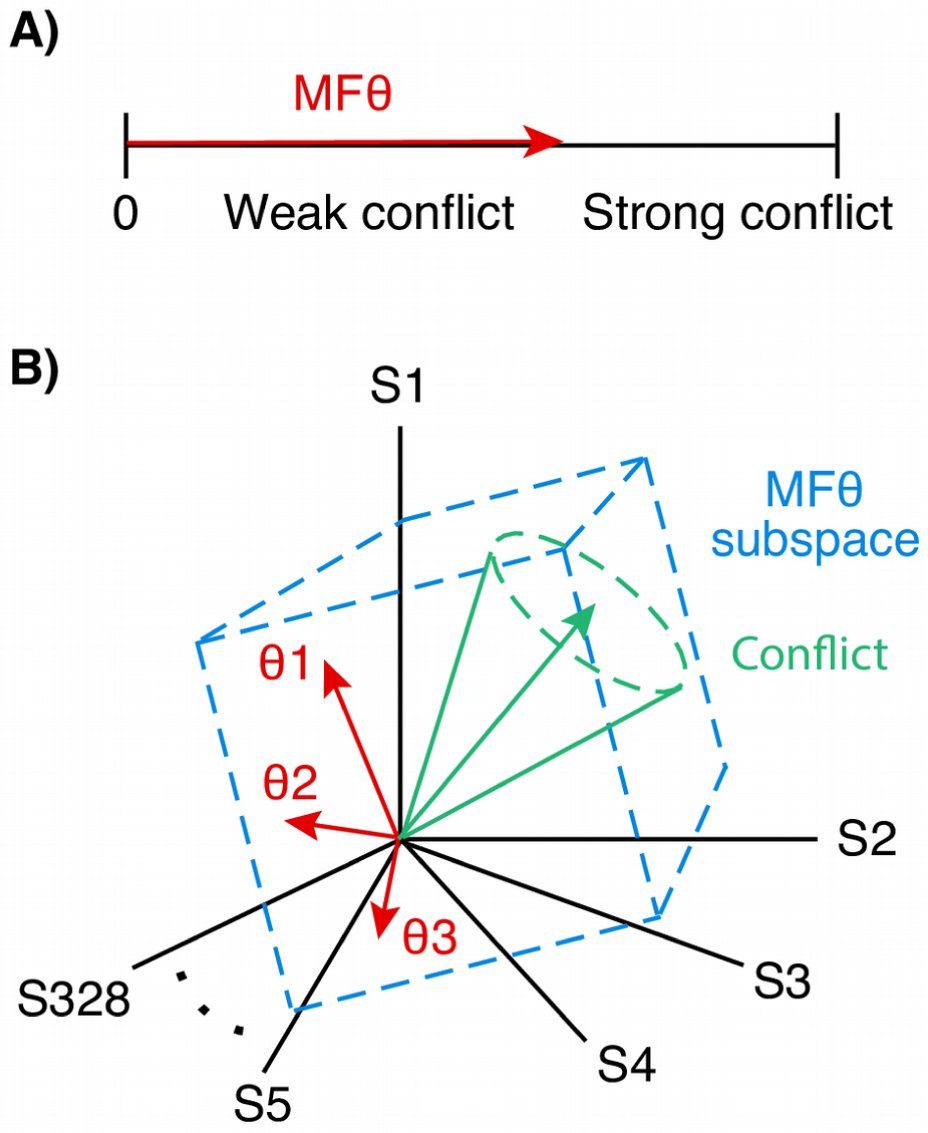
Conceptual visualization of midfrontal theta as a unidimensional vs. a multidimensional phenomenon. **A)** In the classical view, response conflict theta is a singular phenomenon that varies in magnitude with the level of conflict. **B)** Our results underpin a characterization of midfrontal conflict theta as occupying a multidimensional subspace within sensor space (sensors S1 through S328). The independent variability of the basis vectors (shown as theta components θ1 through θ3) implies the existence of multiple independent generators of midfrontal theta, a subset of which are conflict-relevant. As conflict increases, theta activity increases, and the signal state moves further away from the origin.

GED, the feature-guided source separation method used here, has several advantages over “blind” source separation methods like PCA and ICA. Whereas PCA and ICA optimize for sources that best describe the entire dataset, GED optimizes for sources that maximally separate two specified datasets, effectively maximizing the signal-to-noise ratio (de Cheveigné and Parra, 2014). Thus, given a justified a priori selection of data features of interest, GED can achieve higher sensitivity to relevant data features. This sensitivity may explain discrepancies between the current work and the study by Töllner et al. (2017), which used ICA to identify midfrontal theta generators. Töllner et al. described two frontal midline theta clusters spanning multiple subjects. One cluster was localized to MFC and generated more theta-frequency activity in the presence of conflict; the other was localized to the more anterior mPFC and was unmodulated by conflict. Importantly, these clusters manifested with dissociable EEG topographies. We here report multiple components per subject with highly similar EEG topographies (e.g., Figure 3), selected using an FCz-centric template that corresponded closely to the MFC cluster topography found by Töllner et al., Furthermore, Figure 6C shows that a majority of the components we identified generated more theta-frequency activity in the presence of conflict. In contrast, Töllner et al. reported at most a single IC per subject that was modulated by conflict. Given the sensitivity of GED as compared to ICA, the single IC identified by Töllner et al. may itself be a conglomeration of multiple functionally similar, though linearly dissociable, conflict-related theta sources.

The functional and theoretical implications of our findings warrant further consideration. Components were characterized by linear independence, low synchrony, low temporal correlations, and insensitivity of causal coupling to task features, all providing evidence against computational cooperation. Ergo, the generators are likely to perform independent computations. We can thus rule out that these sources reflect response options which compete with one another through mutual inhibition, one of the prevailing theoretical accounts of response conflict processing (Botvinick et al., 2001; Yeung et al., 2004).

What, then, may be the use of these parallel independent computations? In one case, the computations may be functionally equivalent to one another, suggesting the theta generators harness comparable microcircuitry. It has been proposed that response conflict detection and signaling is achieved by specialized theta-resonant microcircuitry within MFC (Cohen, 2014a). Indeed, the presence of multiple sources of conflict theta could reflect the parallel recruitment of assemblies throughout a general neural substrate that is suited to conflict detection. Such parallelism might serve purposes along the lines of consensus or redundancy processing, or the concurrent processing of differing afferent inputs.

In other cases, the different generators may implement different computations. It is possible that response conflict appears as a single phenomenon at the cognitive/behavioral level, but is implemented by a functionally diverse range of circuits operating in the theta range. Indeed, response conflict is associated with activation in a range of subregions along the medial wall of the prefrontal cortex (Ridderinkhof et al., 2004).

A third explanation may be that response conflict is implemented through the parallel recruitment of brain networks. In this scenario, each component is driven not by a single anatomical source, but by a network of sources distributed throughout cortical (and possibly subcortical) circuits. This explanation would account for the more lateral MEG findings (Figures 1D and 3D). Distinct extra-MFC networks, making distinct contributions to the task (potentially related to working memory, attention, motor control, etc.) could appear as distinct components.

From the previous two points, it follows that the midfrontal theta dimensions identified here may be related to different aspects of the conflict task. This possibility is consistent with differences in neural dynamics being observed for stimulus vs. response conflict (Nigbur et al., 2012), the modality in which the stimuli are presented (Donohue et al., 2012; Castro et al., 2018), the complexity of the response mapping (Donohue et al., 2016), and the speed-accuracy tradeoff (Pastötter et al., 2012). Furthermore, the exact localization of conflict theta often depends on the task and on stimulus modality (Ridderinkhof et al., 2004). It is possible that our dimensions originate from spatially distinct midfrontal structures that cannot be distinguished with the spatial resolution of M/EEG, but that can be accessed through linear decomposition. By systematically varying or isolating task features, future studies may be able to determine which dimensions of midfrontal theta, if any, covary with different task or conflict requirements.

In interpreting and generalizing the results of this study, some limitations should be kept in mind. First, statistically separable sources do not equate to anatomically local generators of activity. A statistical source is at its core a spatial filter, the identification of which may have been driven by a single anatomical generator or by distributed anatomical generators that were strongly coupled. As such, the identified components clearly reflected a combination of electrodes and sensors with strong correlational patterns, but are not necessarily driven by a single spatially restricted dipole. Nonetheless, the decomposition of midfrontal theta into multiple components demonstrates that conflict-related midfrontal theta is distributed in a multidimensional subspace, as opposed to being captured by a single statistical source (see also de Cheveigné and Parra, 2014).

Furthermore, our analyses necessitated some design choices. We selected the theta frequency band (5-7 Hz) as our frequency range of interest, based on strong a priori expectations about the manifestation of response conflict. This selection means that our analysis could not capture potentially conflict-relevant sources in other frequencies. Likewise, our explicit template-based selection of midfrontal sources may have excluded relevant sources with different topographical manifestations, such as those that are driven by parietal, visual, or motor sources. However, the selection was applied only to the EEG topographies, while the MEG topographies were left unconstrained. It is thus possible that we captured spatially distributed networks of which the non-midfrontal dipoles were obscured to EEG.

In this work, we have challenged the notion that midfrontal conflict theta is a singular phenomenon. Advanced source separation methods on a multimodal dataset have revealed the presence of multiple theta sources in each subject. Thus, conflict theta, which appears as a single process without close inspection, consists of the composite activity of multiple dissociable theta generators that are likely to implement independent computations. This conceptualization of response conflict theta as a multidimensional phenomenon allows for new functional accounts of conflict processing. Future studies should validate the existence and investigate the nature of these theta sources in more detail, for example in electrophysiological data recorded at smaller spatial scales.

## Acknowledgments

MBZ and MXC are funded by ERC-StG 638589

The authors declare no competing financial interests.

